# Effects of predictability and error on corticospinal excitability during an electronic prediction game

**DOI:** 10.64898/2026.08.30.748085

**Authors:** Victor Hugo Moraes, Bia L. Ramalho, Priscila Azevedo, Aline Duarte, Jesus E. Garcia, Paulo Cabral-Passos, Claudia D. Vargas

## Abstract

**Background:** Detecting environmental regularities is a sophisticated cognitive skill, but its neural basis remains unclear. We tested whether corticospinal excitability tracks stimulus predictability, prediction errors, and task progression while the participants played the Goalkeeper Game.

**Methods:** Sixteen right-handed males (24.4 ± 6.8 years) saved penalty kicks using index, middle, or ring fingers (left, center, right). Kick sequences were generated by a context tree model, in which contexts represent the minimum recent history required to predict the next event. Participants completed 1,200 trials (6 blocks × 200 trials each). Transcranial magnetic stimulation (TMS) delivered 400 ms before the go signal elicited motor evoked potentials (MEPs) in the first dorsal interosseous (FDI) and flexor digitorum superficialis (FDS) muscles during blocks 2, 4, and 6. Contexts were classified as unpredictable (1, 10) or predictable (2, 20, 00); prediction errors were defined for non-deterministic transitions after context 1. Log-MEPs were analyzed using mixed-effects models.

**Results:** In FDI, a Predictability × Previous-Error × Block interaction emerged (F(2,8997)=6.89, p=.001), confined to the final block. Predictability increased MEP amplitudes after successful transitions (+6.3%; p=0.006) but decreased MEP amplitudes after failed transitions (−7.0%; p=0.007). For FDS, unpredictable trials elicited larger MEPs (+7.2%; p=0.002), and MEPs increased from block 2 to 6 (+16.8%; p=0.003).

**Conclusions:** FDS MEP responses were broadly associated with uncertainty and task progression, whereas FDI MEP responses showed context-specific, error-dependent modulation after extended exposure, suggesting fine-grained predictive coding in goal-executing effectors.

## Introduction

The ability to identify statistical patterns in the environment is essential for anticipating future events and optimizing motor responses [1,2]. In motor control, this ability is particularly relevant because movement planning requires decisions under sensory, motor, and task uncertainty, especially when rapid actions are needed [3]. When action selection relies primarily on sensory information available at the moment of response, processing delays and the speed–accuracy trade-off may constrain performance [3]. Thus, learning the probabilistic structure of a sequence of events can facilitate motor preparation and reduce uncertainty about future actions.

A classic approach to investigating sequential pattern learning is the serial reaction time task [4,5]. In deterministic sequences, performance improves as reaction times progressively decrease as the stimulus order is learned [4]. In probabilistic sequences, however, the participant must learn both the order of events and the transition probabilities between them [5]. This allows us to investigate how the brain handles varying degrees of predictability and uncertainty during response preparation.

Context tree models provide a formal framework for generating and studying probabilistic sequences [6,7]. In this model, the probability of the next event depends on a finite set of previous events, known as *context* [8]. Thus, different contexts show varying levels of predictability, allowing comparison of responses to more or less predictable transitions. Recent studies have applied this approach to the Goalkeeper Game, in which participants anticipate the direction of penalty kicks generated by a probabilistic sequence [9,10]. These studies show that participants reduce their response times (RTs) as they gain experience, suggesting learning of the task’s probabilistic structure. Although the behavioral effects of probabilistic learning are relatively well described, their neurophysiological correlates remain less understood [11]. A particularly relevant measure is corticospinal excitability, usually estimated by the amplitude of motor evoked potentials (MEPs) obtained after transcranial magnetic stimulation (TMS), which reflects the momentary excitability of the primary motor cortex, corticospinal tract, and spinal circuits [12]. Contemporary evidence indicates that corticospinal excitability is modulated by preparation, action selection, uncertainty, and surprise, suggesting that the motor system participates in broader stages of predictive action control [13,14].

Despite this, it remains unclear how statistical regularities in event sequences modulate corticospinal excitability. In particular, it remains unknown whether this excitability differentiates between predictable and unpredictable contexts, whether it is influenced by the outcome of the previous prediction, and whether these factors interact over the course of task exposure. Furthermore, to the best of our knowledge, no study has investigated whether intrinsic and extrinsic hand muscles exhibit distinct patterns of corticospinal modulation during a probabilistic prediction task. This distinction is relevant because intrinsic muscles, such as the first dorsal interosseous (FDI), are associated with fine motor control and may exhibit greater functional specificity since evidence indicates that it receives denser corticomotoneuronal projections than extrinsic muscles, such as the flexor digitorum superficialis (FDS), which participate in more general motor synergies [15].

Therefore, the present study aimed to investigate whether contextual predictability, the outcome of prior predictions, and task progression modulate RTs and corticospinal excitability of the FDI and FDS muscles while participants performed the Goalkeeper Game, a prediction task based on a probabilistic sequence generated by a context tree model. We hypothesized that contextual predictability and progressive exposure to the sequence would modulate corticospinal excitability. Furthermore, we postulated that the modulation in the intrinsic hand muscle FDI would more closely track the fine-grained probabilistic structure of the kicker’s sequence than in the FDS.

## Methods

### Participants

The ethics committee of the Institute of Neurology Deolindo Couto, Federal University of Rio de Janeiro (INDC-UFRJ), Brazil, approved the experimental procedures (CAAE:62924022.8.0000.5261). Sixteen right-handed male volunteers (24.4 ± 6.8 years old; handedness 80.3 ± 16.2) were recruited [16], and a TMS safety questionnaire was administered before data collection [17]. The protocol consisted of applying single TMS pulses during motor preparation while participants played the Goalkeeper Game.

### The Goalkeeper Game An electronic prediction game

The Goalkeeper Game is an electronic prediction game in which the player acts as a soccer goalkeeper, aiming to save as many penalty kicks as possible (https://game.numec.prp.usp.br/; v180629a). The volunteer estimates where the penalty taker will kick by pressing the left (→), down (↓), or right (→) arrow keys, making the goalkeeper save to the left, center, or right, respectively. Volunteers responded only after the go signal (red arrows on the screen) and were instructed to defend as many kicks as possible.

In the game, a context tree model [9,10] defined the sequence of penalty kicks, based on the premise that the brain identifies, learns, and stores a finite amount of past information to predict subsequent events. Kicks to the left, center, and right are represented by the symbols 0, 1, and 2, respectively, and five contexts (*w*) were used: 00, 10, 20, 1, and 2 (Fig. 1A). Some contexts are deterministic (e.g., two left kicks are always followed by a center kick, so *p*(1 | 00) = 1), whereas context 1 is not (*p* < 1): given context 1, symbol 0 occurs with 30% probability and symbol 2 with 70%. An example of a generated sequence is 1-2-0-1-0-0-12-0. The model thus generates predictable contexts (2, 20, and 00) and unpredictable ones (1 and 10): context 1 because its successor could be symbol 2 (more probable) or 0 (less probable), and context 10 because it occurred in only 30% of the transitions after symbol 1. No information about the structure of the penalty taker’s sequence was given.

**Figure 1:**
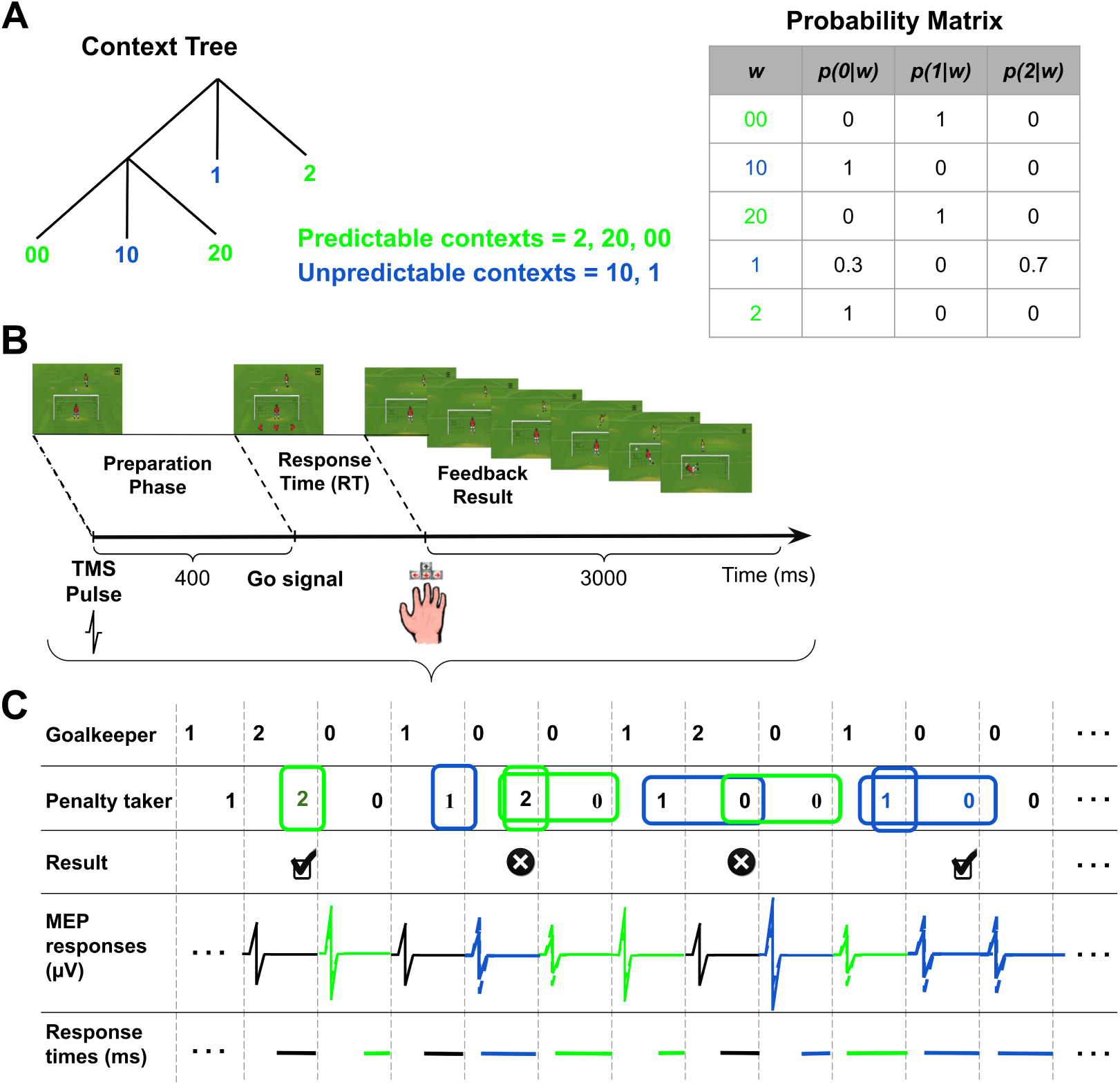
Experimental paradigm and task design. **(A)** Context tree and transition probability matrix used to generate the penalty taker’s sequence. A context (w) is defined as the smallest amount of information required to predict the next event. The probability matrix columns *p*(0|w), *p*(1|w), and *p*(2|w) indicate the probability of occurrence of the symbols 0, 1, or 2, respectively, given each context. The contexts classified as unpredictable (blue) were contexts 1 and 10. The other contexts were classified as predictable (green). **(B)** Temporal evolution of one trial in the game. The TMS pulse was applied 400 ms before the red arrows (go signal), while RTs were recorded as the time between the appearance of the arrows on the screen and the goalkeeper’s response. The feedback animation occurred immediately after the goalkeeper’s response. **(C)** Sequence of trials in the game and grouping of variables based on their predictability. At each trial, the goalkeeper must decide whether to save to the left (0), the center (1), or the right (2). Subsequently, the penalty taker kicks in one of the three directions, depending on the context tree model that governs the sequence. Results of the predictions in non-deterministic transitions were recorded for data analysis. We grouped MEPs and RTs based on context predictability.

At each trial, the TMS pulse was applied during the motor preparation phase [18], 400 ms before the go signal (arrows) appeared on the screen. Following the go signal, the goalkeeper made a choice, and feedback indicated whether that choice was correct (Fig. 1B).

Prediction errors were computed for each non-deterministic transition (context 1) (Fig. 1C): a correct prediction of the symbol following context 1 was a success (✓) and an incorrect one a failure (×). We investigated how contextual predictability (green or blue) and prediction errors modulated MEPs and RTs.

Volunteers were seated comfortably about 114 cm from the screen (LCD Display++, Cambridge Research Systems Ltd), with their hands on an ergonomic keyboard (RB-840, Cedrus Co.) (Fig. 2A, B). The protocol was performed in a single ∼3-hour visit, including preparation and data collection, and consisted of 1 familiarization block (12 trials) and 6 experimental blocks (200 trials each; ∼16 min per block). Single TMS pulses were applied in blocks 2, 4, and 6, while blocks 1, 3, and 5 were performed without TMS, with a 1–2 min interval between blocks to promote recovery and maintain engagement. Due to coil overheating, pulses were applied only after the initial learning block, alternating between stimulated and non-stimulated blocks.

**Figure 2:**
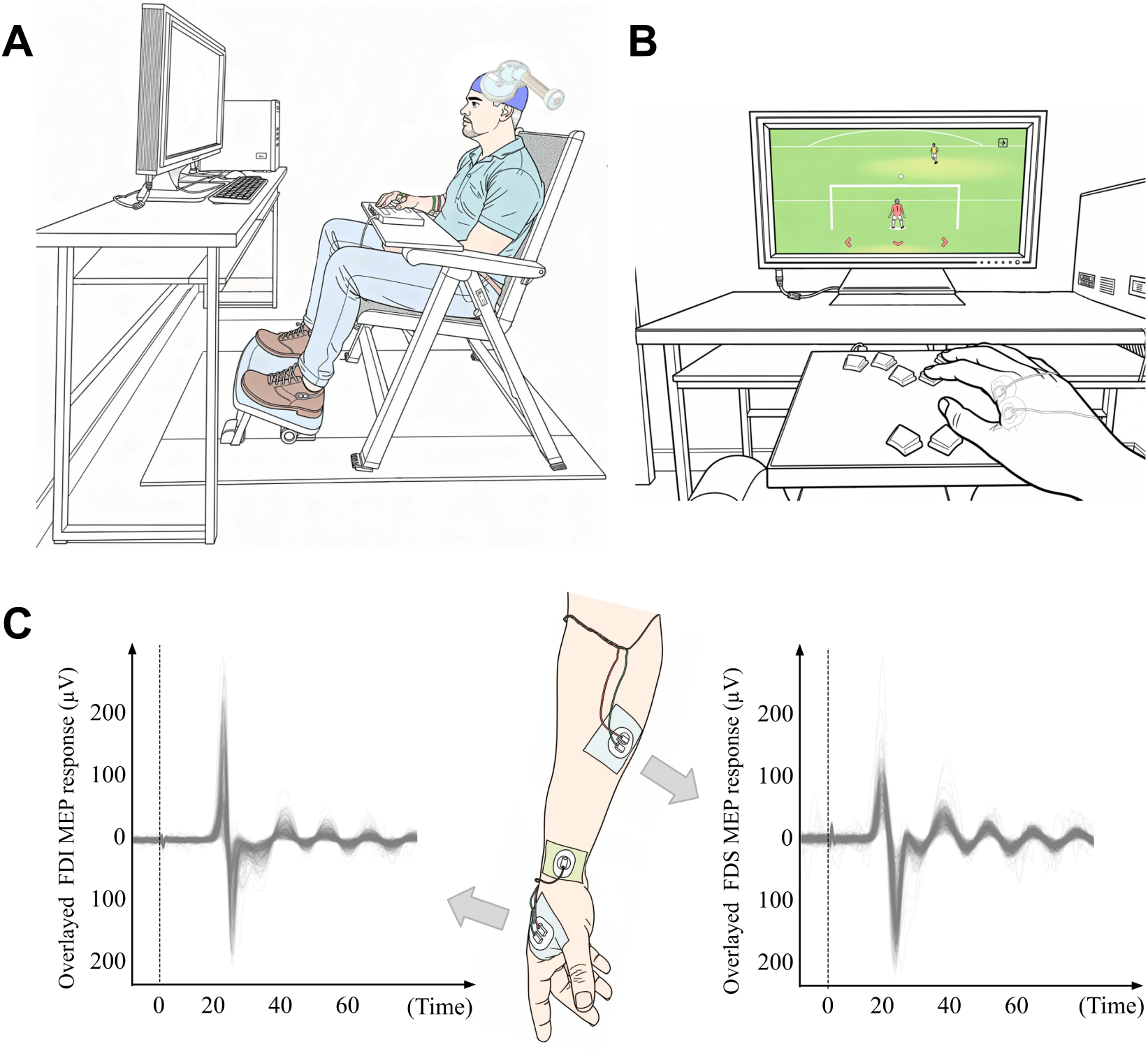
Experimental setup and TMS protocol. **(A)** Position of the goalkeeper and acquisition of MEPs through surface electromyography (sEMG). Volunteers sat comfortably about 114 cm from a monitor, with their right (dominant) hand on a directional pad to interact with the Goalkeeper Game. The TMS coil was kept over the hotspot of the FDI muscle, oriented at approximately 45 degrees (relative to the sagittal plane). **(B)** The volunteer’s view while playing as a goalkeeper during the game. The index, middle, and ring fingers were placed on keyboard keys to select left, center, and right saves, respectively. **(C)** MEPs for FDI (left panel) and FDS (right panel) from a representative goalkeeper were superimposed for visualization. The vertical dotted line represents the pulse time (time 0).

### TMS protocol

A Magstim 200² stimulator (Magstim Co., UK) with a D70² figure-of-eight coil was used for focal stimulation. Coil placement was maintained using the open-source InVesalius Navigator software [19], a Polhemus Patriot tracker (Polhemus Co., USA), and an MNI template head model, with a maximum target-localization error of 3 mm. Surface electromyographic (sEMG) activity was recorded from the FDI and FDS muscles (Fig. 2C) using an ActiCHamp amplifier and BrainVision Recorder software (Brain Products, Germany); MEPs were derived from these recordings. Ambu Neuroline 715 passive Ag/AgCl surface electrodes (71512-K/C/12) were connected to an analog differential DC preamplifier (gain: 100; ±3% tolerance). Signals were digitized at 5 kHz and exported for offline processing.

The FDI hotspot was located within a 12-point matrix (1 cm spacing) drawn on the cap around a point at 20% of the Cz–left tragus distance. Single monophasic pulses (40% intensity, coil ∼45° to the median plane, increased in 2% steps when needed) were delivered to each point at 4–8 s intervals; the site eliciting the largest mean FDI MEPs over 5 pulses was defined as the hotspot and recorded by the neuronavigation system for replication throughout the experiment. Resting motor threshold (rMT) was determined from the hotspot-defining intensity using the ML-PEST algorithm (MTAT 2.0 software [20]). MEPs were then recorded at 120% rMT with ∼5 s inter-stimulus intervals.

### Data processing

RTs were calculated as the interval between the go signal and the volunteer’s button press. The raw sEMG signal was filtered with a 60 Hz notch filter and a 20–500 Hz Butterworth (2nd-order) bandpass filter to attenuate line noise and spurious spectral components. Using the TMS pulse trigger in the sEMG signal, 50-ms windows (10–60 ms after the trigger) were extracted to quantify MEP peak-to-peak amplitude (max−min, *µ*V) in the FDI and FDS muscles. RTs greater than 1.5 s and MEPs with prestimulus baseline myoelectric activity (RMS over the 500 ms preceding stimulation) exceeding 2 standard deviations above the participant-specific mean were excluded from analysis. Data processing was conducted in Python 3.11 (Python Software Foundation), mainly using the MNE and Pandas libraries.

### Statistical analysis

Statistical analyses were conducted in R (v4.5.2) using the rstatix, lme4, lmerTest, emmeans, and effectsize packages. Log-transformed MEPs and RTs were analyzed with linear mixedeffects models (LMMs; REML estimation, Satterthwaite approximation), with fixed effects of Predictability (Predictable/Unpredictable), Previous-Error (Success/Failure), and Block (2, 4, 6), their two- and three-way interactions, and trial position as a covariate. The maximal model included by-participant random intercepts and uncorrelated random slopes for all predictors (convergence and non-singularity confirmed). For RTs, however, the maximal model produced a singular fit, so a parsimonious model retaining only by-participant random intercepts (same fixed-effects structure) was used.

Fixed effects were tested via Type III ANOVA, with partial eta-squared (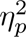) as effect size and model fit summarized by marginal and conditional R². Significant interactions were followed by estimated marginal means (EMMs) and pairwise contrasts, back-transformed to the response scale (µV for MEPs, ms for RTs) and adjusted with the Benjamini–Hochberg procedure. Percentage differences were derived from contrast ratios as (ratio 1) × 100, relative to baseline.

To verify that findings did not depend on specific modeling choices, a complementary repeated-measures ANOVA (RM-ANOVA) was performed on cell-aggregated MEP/RT measures (mean per participant × condition), with Greenhouse–Geisser correction for sphericity violations and 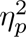 as effect size. Convergence between LMM and RM-ANOVA results was taken as evidence of robustness.

Success rate across blocks was compared using a Kruskal–Wallis test, followed by Dunn’s post-hoc tests with Benjamini–Hochberg correction; effect size was estimated as eta-squared (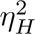). Code for data processing and statistical analysis is available on GitHub (https://github.com/moraesvictorhugo/TMS-GoalkeeperGame).

## Results

MEP amplitudes in the FDI and FDS muscles, together with RTs from the Goalkeeper Game, were analyzed to test for modulation by contextual predictability, prediction error, and temporal evolution across the experimental session.

For FDI, log-transformed MEP amplitudes were modeled with a linear mixed-effects model (REML, Satterthwaite df) including the Predictability × Previous-Error × Block interaction, trial position, and uncorrelated by-participant random intercepts and slopes for all predictors (9082 observations, 16 participants; non-singular). No main effects reached significance (Predictability: F(1, 15.4) = 0.06, *p* = 0.805, 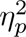 = 0.004; Previous-Error: F(1, 14.9) = 0.65, *p* = 0.431, 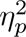 = 0.04; Block: F(2, 15.1) = 0.32, *p* = 0.731, 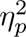 = 0.04; trial position: F(1, 15.0) = 0.81, *p* = 0.381, 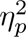 = 0.05). Instead, effects emerged through the Predictability × Previous-Error (F(1, 8997) = 8.86, *p* = 0.003, 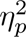 = 0.001) and three-way (F(2, 8997) = 6.89, *p* = 0.001, 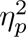 = 0.002) interactions, confined to Block 6 (all contrasts *p* > 0.20 in Blocks 2 and 4). There, the predictability effect reversed with prior outcome: after success, predictable trials elicited larger MEPs than unpredictable ones (+6.3%; ratio = 1.063, z = 2.77, *p* = 0.006), whereas after failure the effect inverted (−7.0%; ratio = 0.930, z = −2.68, *p* = 0.007). Equivalently, success increased excitability relative to failure in predictable trials (+6.2%; ratio = 1.062, z = 2.37, *p* = 0.018) but decreased it in unpredictable ones (−7.0%; ratio = 0.930, z = −2.81, *p* = 0.005). This crossover suggests corticospinal excitability tracks the congruence between contextual predictability and prior outcome after extended task exposure (Block 6; Fig. 3). A sensitivity RM-ANOVA on trial-averaged data confirmed the three-way interaction (Supplementary Results: Table 5).

**Figure 3:**
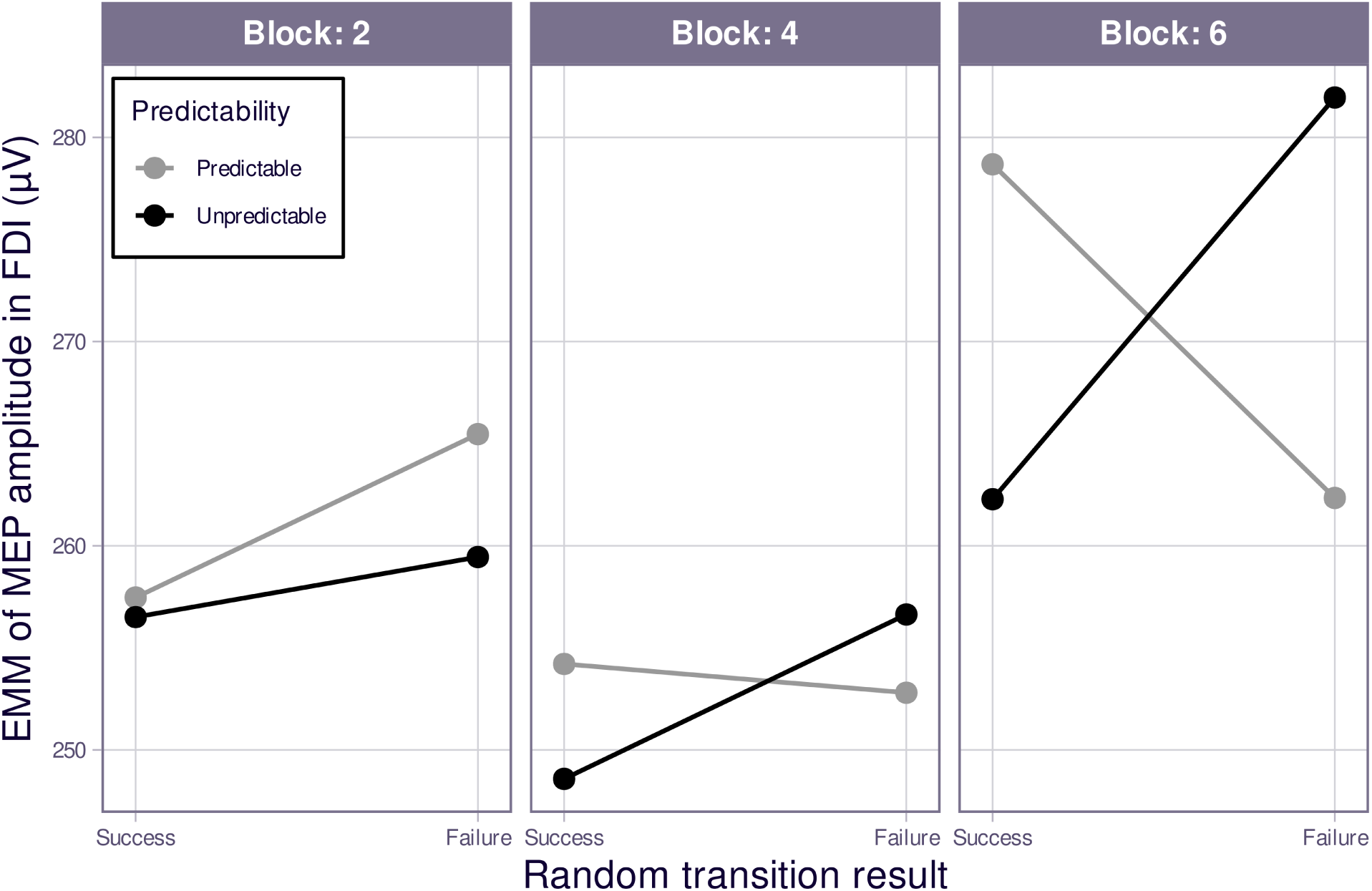
Effects of block (2, 4, 6), previous prediction outcome in non-deterministic transitions (failure, success), and contextual predictability (predictable, unpredictable) on MEP amplitude in the FDI muscle. The y-axis shows estimated marginal means (EMMs) of the MEP amplitude (µV) derived from the linear mixed-effects model (LMM). The x-axis is organized in two hierarchical levels: block at the outer level (Block 2, Block 4, Block 6) and previous prediction outcome at the inner level (failure, success). Line colors represent context predictability (gray: predictable; black: unpredictable). n = 16.

For the FDS, log-transformed MEP amplitudes were modeled with the same mixed-effects structure (REML, Satterthwaite degrees of freedom; maximal Predictability × Previous-Error × Block interaction, trial position within block, and uncorrelated random intercepts and slopes per volunteer for all four predictors; 9341 observations, 16 participants). The model converged normally and was not singular. In contrast to the FDI, the FDS showed two robust main effects rather than an interaction (Fig. 4). MEPs were 7.2% larger on unpredictable than predictable trials (F(1, 15.3) = 14.79, *p* = 0.002, 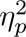 = 0.49; ratio = 1.072) and increased gradually across the session (F(2, 15.2) = 7.54, *p* = 0.005, 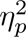 = 0.50), rising 16.8% from Block 2 to Block 6 (ratio = 1.168, z = 3.32, *p* = 0.003). Previous-Error had no main effect (F(1, 15.0) = 0.99, *p* = 0.336, 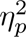 = 0.06), and trial position showed only a non-significant trend (F(1, 15.0) = 3.34, *p* = 0.088, 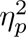 = 0.18). None of the interactions reached significance—including the Predictability × Previous-Error (F(1, 9270) = 2.84, *p* = 0.092, 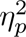 < 0.001) and three-way (F(2, 9262) = 0.04, *p* = 0.956, 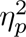 < 0.001) terms that had driven the FDI effects. A sensitivity RM-ANOVA on trial-averaged data reproduced the main effects of predictability and block (Supplementary Results: Table 10).

**Figure 4:**
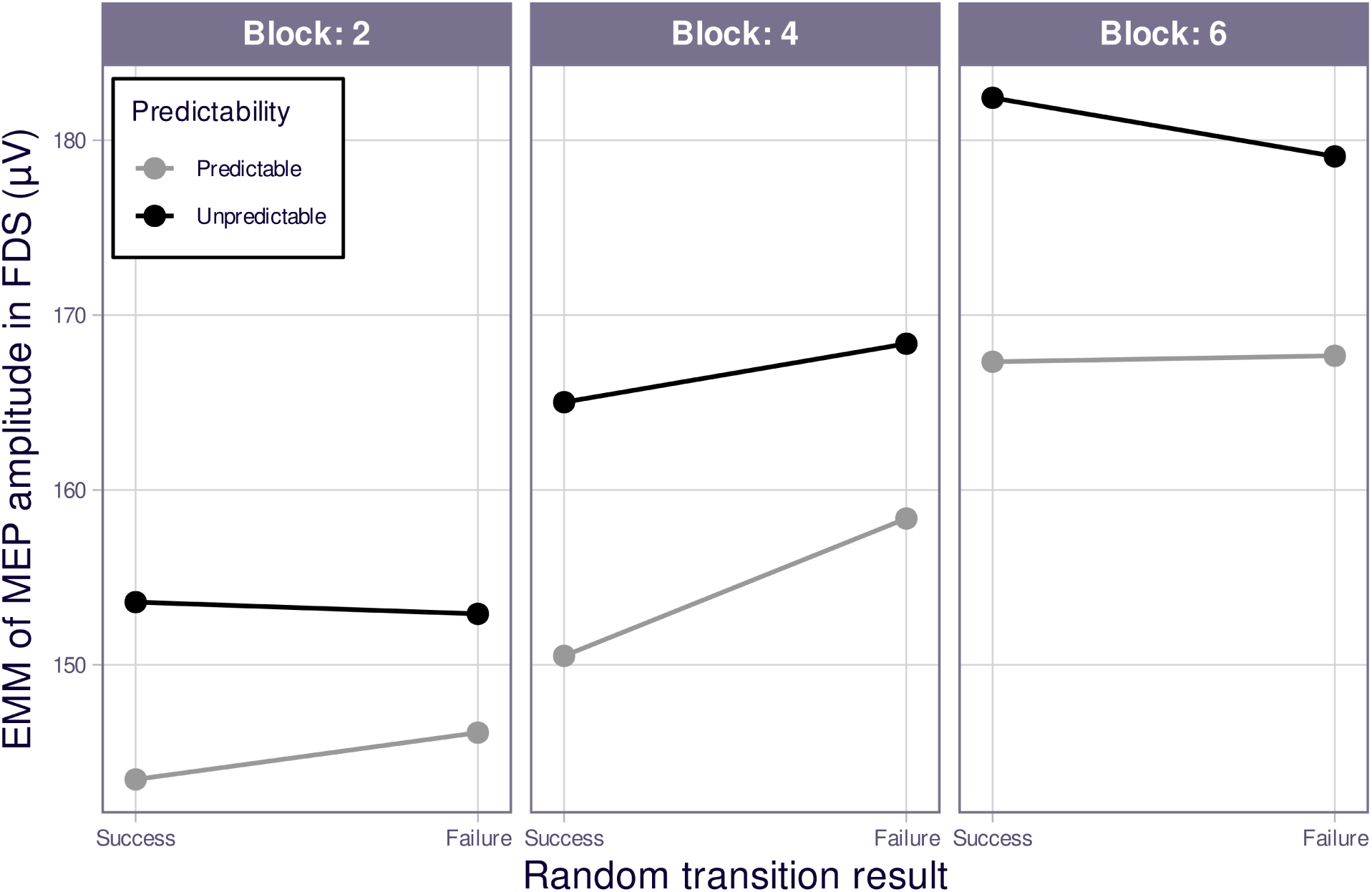
Effects of block (2, 4, 6), previous prediction outcome in non-deterministic transitions (failure, success), and contextual predictability (predictable, unpredictable) on MEP amplitude in the FDS muscle. The y-axis shows estimated marginal means (EMMs) of the MEP amplitude (µV) derived from the linear mixed-effects model (LMM). The x-axis is organized in two hierarchical levels: block at the outer level (Block 2, Block 4, Block 6) and previous prediction outcome at the inner level (failure, success). Line colors represent context predictability (gray: predictable; black: unpredictable). n = 16.

Across the game, RTs decreased progressively (Block 2 → Block 6: 537 → 415 ms; F(2, 9194) = 301.40, *p* < 0.001, 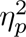 = 0.06; Fig. S1) and success rates increased from Block 1 onward (Block 1: 0.68 ± 0.14 vs. Block 6: 0.85 ± 0.03; H(5) = 25.7, *p* < 0.001, 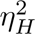 = 0.229; Fig. S2). Full analyses are provided in the Supplementary Results.

## Discussion

This study aimed to evaluate whether contextual predictability or prediction errors in a predictive electronic game would modulate corticospinal excitability. Volunteers acted as a goalkeeper and had to save as many penalty kicks as possible, with the penalty taker’s sequence defined by a context tree model in which kicks depended on both past context and its transition probabilities [10]. The results showed that: (1) FDI MEPs showed an interaction between the previous prediction outcome (in non-deterministic transitions) and contextual predictability in the final stage of learning, with larger MEPs for previous failure + unpredictable context and previous success + predictable context; (2) FDS MEPs increased across learning and were consistently larger in unpredictable than in predictable contexts, regardless of the previous prediction outcome; and (3) RTs decreased and success rate increased as learning progressed.

### Effect of context predictability on MEPs

Based on previous studies, we hypothesized that contextual predictability would modulate corticospinal excitability. As highlighted by Bestmann and colleagues [12], corticospinal excitability reflects the state of M1 and the corticospinal tract at a given moment, since M1 receives excitatory and inhibitory inputs from different brain networks [12,21]. We therefore hypothesized that contextual predictability would affect MEP amplitude in both the FDI and FDS. As shown in Figures 3 and 4, predictability modulated MEPs in both muscles, but differently: as a main effect in the FDS (larger MEPs on unpredictable trials), and only in interaction with the previous outcome in the FDI. Because the pulse was delivered 400 ms before the go signal, these modulations likely resulted from a combination of motor preparation [22] and cognitive-motor processes engaged in processing past information, i.e., contextual predictability [12,13].

Evidence that probabilistic context modulates corticospinal excitability remains limited. Bestmann et al. [13] showed that MEP amplitudes during action preparation were sensitive to cue validity, block-level predictability, and trial-by-trial surprise, being larger when the upcoming action was more predictable, when cues were valid, and when the previous trial was less surprising. Our FDI results partially align with these findings: after successful predictions, FDI MEPs were greater in predictable than unpredictable contexts, particularly after extended exposure to the task. For the FDS, however, the findings appear to differ, showing increased excitability in unpredictable contexts. Some methodological differences may account for this discrepancy. First, in their work, predictability was defined by explicit cue–action associations and block-level probabilities, whereas in the present task it emerged from sequential statistical structure. Second, they delivered TMS 200 ms before the go signal, whereas stimulation here occurred 400 ms before it; thus, MEPs may have sampled different stages of decision-making and motor preparation. Together, these findings suggest that corticospinal excitability, especially in intrinsic hand muscles directly involved in the selected action, is shaped not only by contextual predictability and surprise, but also by the recent history of prediction outcomes.

### Crossover interaction between prediction error and context predictability modulates FDI MEPs

Our analysis showed that, after prolonged exposure to the Goalkeeper Game (Block 6, ∼1,000 plays), FDI corticospinal excitability was modulated by the interaction between contextual predictability and the outcome of the preceding non-deterministic transition, rather than by either factor alone (Fig. 3). The pattern took the form of a crossover: after a successful prediction, MEPs were larger in predictable than in unpredictable contexts (+6.3%), whereas after a failure this pattern reversed, with larger MEPs in unpredictable contexts (predictable trials: −7.0%). Equivalently, previous success enhanced excitability relative to failure under predictable contexts but suppressed it under unpredictable ones. Although a directionally consistent trend was already visible in Block 4, it did not reach significance (all *p* > .20) and became reliable only in Block 6. These findings suggest that the context-dependent modulation of corticospinal excitability requires extended task experience. For the FDS, by contrast, there was no interaction, only main effects of increased MEPs across learning and consistently larger MEPs in unpredictable than in predictable contexts, regardless of the previous prediction outcome. These results raise a fundamental question: why was this modulation, dependent on contextual predictability and prediction error, evident only in the FDI and not the FDS?

Previous studies have investigated how errors affect corticospinal excitability in intrinsic and extrinsic hand muscles. In an Eriksen Flanker Task, Amengual et al. [23] had participants respond with the index or middle finger to compatible (HHHHH/SSSSS) or incompatible (HHSHH/SSHSS) letter patterns, delivering single TMS pulses over M1 ipsilateral to the active hand at 150, 300, or 450 ms after correct or erroneous responses. FDI MEP amplitude increased 450 ms post-error; given the inverse relationship between the two M1 hemispheres, the authors inferred decreased excitability of the active motor cortex following errors, potentially preventing premature or repeated erroneous responses. Suzuki et al. [24] examined corticospinal excitability during trial-and-error decision-making: colored circles cued different reward probabilities (10–90%), and participants decided whether to perform wrist flexion, receiving a reward or penalty accordingly. A TMS pulse was applied 1 s after feedback at the midpoint between the centers of gravity of the flexor carpi radialis (agonist) and extensor carpi radialis (antagonist), and agonist MEP amplitudes were significantly larger under penalty than reward. Together, these findings suggest errors can modulate corticospinal excitability in both intrinsic and extrinsic muscles, either increasing or decreasing it. However, none of these protocols recorded intrinsic and extrinsic muscle activity simultaneously, precluding a direct comparison.

This study therefore supports an alternative hypothesis: that corticospinal excitability in an intrinsic hand muscle—given its greater specialization as an effector—can be modulated more precisely. Anatomically, three factors support this. First, the innervation ratio (muscle fibers per motor neuron) of intrinsic hand muscles is lower than that of extrinsic muscles [25,26]; because smaller motor units allow more gradual force increments, force gradation is finely tuned, rendering changes in corticospinal excitability more readily detectable. Second, the FDI has a higher density of monosynaptic corticomotoneuronal projections, so M1 output reaches FDI motor neurons more directly than those of the FDS, with less interference from local spinal circuits [27]. Third, the cortical representations of the FDI and FDS reflect distinct functional synergies in M1: the FDI overlaps more with other intrinsic hand muscles (e.g., abductor pollicis brevis and flexor pollicis brevis), whereas the FDS overlaps substantially with forearm muscles (e.g., flexor carpi radialis, FCR) [28], an organization that may have shaped the task-specific modulation observed. Moreover, MEPs recorded over the forearm and attributed to the FDS may reflect the summation of myoelectric signals from multiple muscles, such as the FCR and palmaris longus. This well-documented crosstalk [29,30] may compromise FDS signal specificity and, consequently, the detection of subtle excitability changes. In addition, the hotspot was optimized for the FDI rather than the FDS; future studies could define an FDS-specific hotspot to test whether these findings persist.

Furthermore, evidence from Tomberg and Caramia [31] suggests that the dissociation between FDI and FDS corticospinal excitability may also reflect a task-specific feature. Applying a simple reaction-time task using both subthreshold and suprathreshold TMS over the left hemisphere, delivered 90 ms after the go signal (brief electric pulse delivered to the left thumb), the authors showed that MEP facilitation was selectively directed to the muscle that would actually execute the intended movement. When subjects prepared a right index finger flexion, TMS facilitated motor-evoked responses in the right FDI muscle, while the synergist right flexor digitorum communis (FDC) muscle showed little or no facilitation and, in some trials, was concurrently suppressed. These findings indicate that motor preparation involves selective, task-dependent activation of corticospinal circuitry toward the prime mover muscle required for the specific movement, rather than diffuse facilitation across synergic muscles. Taken together, these factors may help explain why we observed more refined modulation of corticospinal excitability in the FDI than in the FDS.

### Effect of Predictability over RTs and the improvement in Goalkeeper Game performance over time

Our results showed that the predictability of past contexts significantly modulated RTs (Fig. S1). Similarly, Cabral-Passos et al. [10] observed that in predictable contexts, RTs after errors were slower than after correct responses in non-deterministic transitions, but the opposite was found in an unpredictable context. This result was consistent with the cognitive control theory, which posits that errors trigger processes that prevent the repetition of wrong choices [32]. In our data, however, the failure/success effect emerged only in the unpredictable context, and was small in magnitude. This qualitative divergence was likely due to protocol differences: our version of the Goalkeeper Game was modified to comply with a TMS protocol requiring a minimum interstimulus interval of 4 s [33]; accordingly, the feedback phase was extended from the 450 ms used by Cabral-Passos et al. [10] to 3000 ms, likely attenuating the prediction-error effect on RTs. If so, one might speculate that context predictability exerts an even more persistent modulatory effect than prediction errors.

Success rate is a common performance measure in decision-making paradigms [4,34]; here, volunteers showed significantly higher success rates in Blocks 3–6 than in Block 1. The sequence comprised three events (1, X, and 0), where the non-deterministic X was the symbol 2 in 70% of cases and 0 in 30%. Given the periodic occurrence of 1 and 0, worst-case accuracy would not fall below 66%, whereas the last block showed a minimum of 0.81 and a mean of 0.85 (Fig. S2). Previous studies using the Goalkeeper Game and the context-tree model reported a comparable increase in success rate over time [9,10], indicating a general pattern of performance improvement throughout the task. Together, the higher success rate and shorter RTs over time are consistent with the learning effect expected during the game.

## Conclusions

The Goalkeeper Game, combined with TMS, provides a useful framework to probe how statistical learning shapes corticospinal excitability during motor preparation. Theoretically, the results suggest that the corticospinal system integrates contextual predictability, previous action outcomes, and effector-specific motor preparation. Future studies should determine whether the muscle-specific effects observed here persist when using FDS-specific stimulation hotspots, and high-density sEMG to reduce crosstalk. It would also be important to test different stimulation timings, manipulate feedback duration and intertrial interval, include larger and more diverse samples, and assess whether these effects are retained across days. Such approaches may clarify how predictive learning reorganizes corticospinal dynamics and may inform future motor training or rehabilitation protocols based on adaptive prediction.

## Declaration of generative AI and AI-assisted technologies

During the preparation of this work, the authors used Claude Opus 4.8 to improve language clarity and code syntax. After using this tool, the authors reviewed and edited the content and took full responsibility for the publication.

## Credit authorship contribution statement

**Victor Hugo Moraes:** Conceptualization, Methodology, Investigation, Data curation, Formal analysis, Visualization, Writing – original draft, Writing – review & editing. **Bia Ramalho:** Conceptualization, Methodology, Investigation, Validation, Writing – review & editing. **Priscila Azevedo:** Methodology, Investigation, Writing – review & editing. **Aline Duarte:** Methodology, Formal analysis, Validation, Writing – review & editing. **Jesus E. Garcia:** Methodology, Formal analysis, Validation, Writing – review & editing. **Paulo Cabral-Passos:** Methodology, Formal analysis, Validation, Writing – review & editing. **Claudia D. Vargas:** Conceptualization, Methodology, Supervision, Funding acquisition, Resources, Project administration, Validation, Writing – review & editing. All authors contributed to interpreting the results and approved the final version of the manuscript.

## Supplementary Material

### Supplementary Results

#### Linear Mixed-Effects Model Predicting log(MEP) at FDI

Formula:

log_MEP_FDI ∼ Predictability * Error_Prev * Block_Factor + trial_in_block_z + (1 + P_unpred + E_error + B4 + B6 + trial_in_block_z || volunteer) Obs: 9082 | Groups: 16 volunteers | REML: 8575.5 | Singular: No | Convergence: OK

**Table 1.** FDI MEPs LMM Fixed effects.

| Term | Estimate | SE | df | t | p | Sig. |
| --- | --- | --- | --- | --- | --- | --- |
| (Intercept) | 5.551 | 0.217 | 15.0 | 25.631 | 7.95e-14 | *** |
| Pred. Unpredictable | -0.0037 | 0.0223 | 74.8 | -0.168 | 0.867 |  |
| Error_Prev Failure | 0.0306 | 0.0240 | 66.1 | 1.273 | 0.207 |  |
| Block_Factor4 | -0.0127 | 0.0481 | 17.2 | -0.264 | 0.795 |  |
| Block_Factor6 | 0.0792 | 0.0773 | 15.8 | 1.025 | 0.321 |  |
| trial_in_block_z | 0.0231 | 0.0256 | 15.0 | 0.902 | 0.381 |  |
| Pred×Error | -0.0192 | 0.0282 | 8972 | -0.681 | 0.496 |  |
| Pred×Block4 | -0.0187 | 0.0259 | 8992 | -0.719 | 0.472 |  |
| Pred×Block6 | -0.0569 | 0.0259 | 8997 | -2.198 | 0.028 | * |
| Error×Block4 | -0.0362 | 0.0284 | 9022 | -1.275 | 0.202 |  |
| Error×Block6 | -0.0910 | 0.0287 | 9018 | -3.173 | 0.0015 | ** |
| Pred×Error×Block4 | 0.0566 | 0.0408 | 9001 | 1.388 | 0.165 |  |
| Pred×Error×Block6 | 0.1520 | 0.0412 | 8994 | 3.691 | 0.00023 | *** |

<sup>9</sup> *Note.* Significance codes: \*\*\* $p < 0.001$ , \*\* $p < 0.01$ , \* $p < 0.05$ , . $p < 0.10$ .
<sup>10</sup> **Table 2** *Type III ANOVA (Satterthwaite) + Effect Size*

**Table 2.**
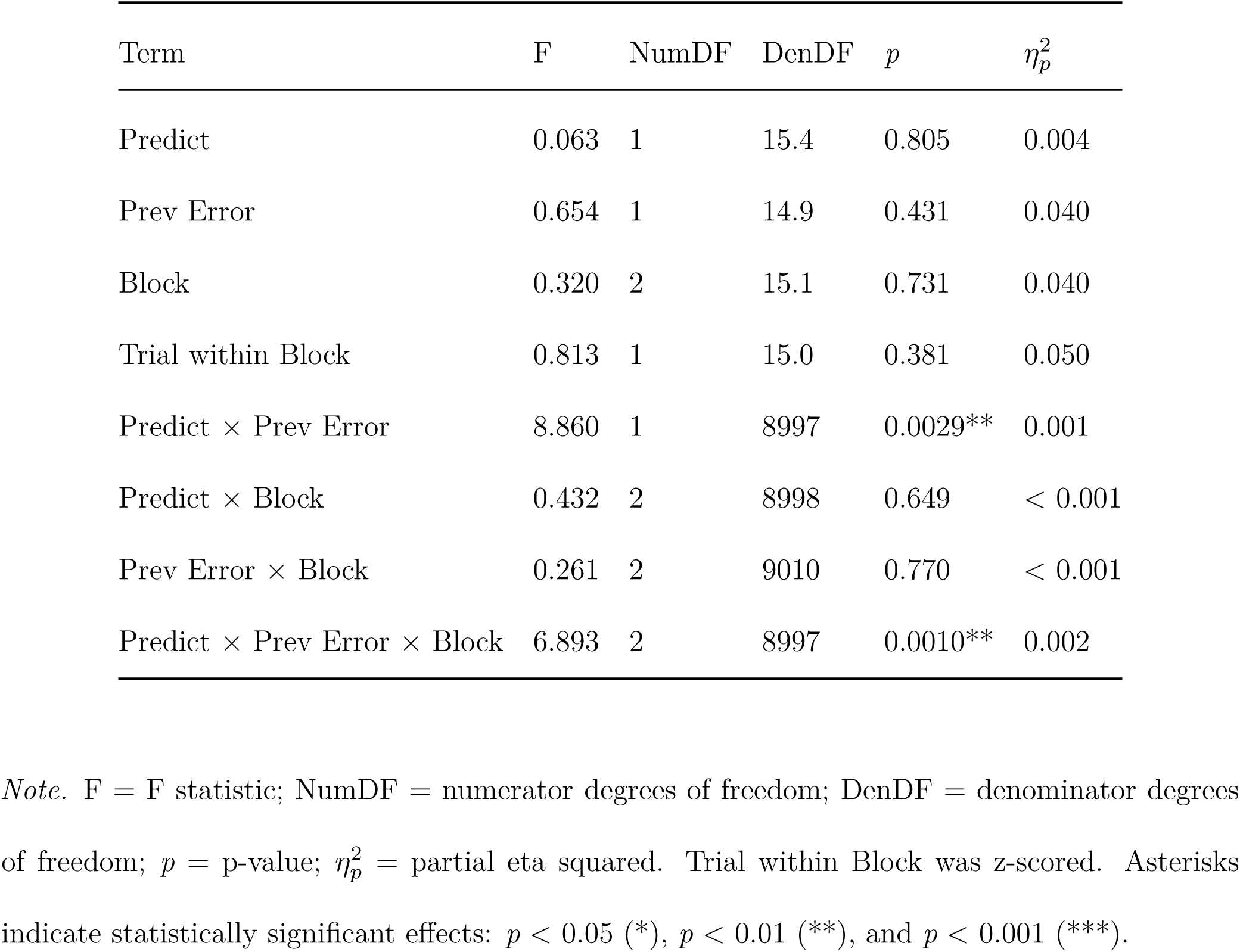
Type III ANOVA (Satterthwaite) + Effect Size.

| Term | F | NumDF | DenDF | p | $\eta_p^2$ |
| --- | --- | --- | --- | --- | --- |
| Predict | 0.063 | 1 | 15.4 | 0.805 | 0.004 |
| Prev Error | 0.654 | 1 | 14.9 | 0.431 | 0.040 |
| Block | 0.320 | 2 | 15.1 | 0.731 | 0.040 |
| Trial within Block | 0.813 | 1 | 15.0 | 0.381 | 0.050 |
| Predict × Prev Error | 8.860 | 1 | 8997 | 0.0029** | 0.001 |
| Predict × Block | 0.432 | 2 | 8998 | 0.649 | < 0.001 |
| Prev Error × Block | 0.261 | 2 | 9010 | 0.770 | < 0.001 |
| Predict × Prev Error × Block | 6.893 | 2 | 8997 | 0.0010** | 0.002 |

<sup>11</sup> *Note.* F = F statistic; NumDF = numerator degrees of freedom; DenDF = denominator degrees <sup>12</sup> of freedom; $p$ = p-value; $\eta_p^2$ = partial eta squared. Trial within Block was z-scored. Asterisks <sup>13</sup> indicate statistically significant effects: $p < 0.05$ (\*), $p < 0.01$ (\*\*), and $p < 0.001$ (\*\*\*).
<sup>14</sup> **Table 3** *Variance Components (random effects)*

**Table 3.** Variance Components (random effects)

| Component | Std.Dev. |
| --- | --- |
| Intercept (volunteer) | 0.865 |
| P_unpred | 0.049 |
| E_error | 0.057 |
| B4 | 0.182 |
| B6 | 0.303 |
| trial_in_block_z | 0.101 |
| Residual | 0.379 |

**Table 4.** Model Fit (R^²^)

| Metric | Value |
| --- | --- |
| Conditional $R^2$ | 0.839 |
| Marginal $R^2$ | 0.002 |

**Table 5.** FDI MEPs analysis — Model’s comparison table.

| Term | Model | F | $p$ | $\eta_p^2$ |
| --- | --- | --- | --- | --- |
| Predictability | Max | 0.063 | 0.805 | 0.004 |
|  | RMA | 0.018 | 0.895 | 0.001 |
| Previous error | Max | 0.654 | 0.431 | 0.042 |
|  | RMA | 0.032 | 0.860 | 0.002 |
| Block | Max | 0.320 | 0.731 | 0.041 |
|  | RMA | 0.443 | 0.598 | 0.029 |
| Trial within block | Max | 0.813 | 0.381 | 0.052 |
|  | RMA | — | — | — |
| Predictability × Previous error | Max | 8.860 | 0.003 | 0.001 |
|  | RMA | 5.447 | 0.034 | 0.266 |
| Predictability × Block | Max | 0.432 | 0.649 | $< 0.001$ |
|  | RMA | 0.784 | 0.424 | 0.050 |
| Previous error × Block | Max | 0.261 | 0.770 | $< 0.001$ |
|  | RMA | 0.397 | 0.635 | 0.026 |
| Predictability × Previous error × Block | Max | 6.893 | 0.001 | 0.002 |
| | RMA | 10.483 | $< 0.001$ | 0.411 |

*Note.* Max = maximal model; RMA = repeated-measures ANOVA; $\eta_p^2$ = partial eta squared.
Dashes indicate that the effect was not included in the RMA model.

#### FDI Cell-aggregate RM-ANOVA

A complementary RM-ANOVA on cell-aggregated MEP amplitudes (192 cells, 16 participants; Greenhouse–Geisser corrected) confirmed this pattern was not model-dependent: no significant main effects (all *p* ≥ 0.598), but significant Predictability × Previous-Error (F(1, 15) = 5.45, *p* = 0.034, 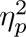 = 0.266) and three-way (F(1.65, 24.82) = 10.48, *p* < 0.001, 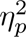 = 0.411) interactions. This agreement across the maximal mixed model and aggregated RM-ANOVA provides converging evidence for the context-dependent modulation of corticospinal excitability reported here.

#### Linear Mixed-Effects Model Predicting log(MEP) at FDS

Formula:

log_MEP_FDS ∼ Predictability * Error_Prev * Block_Factor + trial_in_block_z + (1 + P_unpred + E_error + B4 + B6 + trial_in_block_z || volunteer)

Obs: 9341 | Groups: 16 volunteers | REML: 6403.1 | Singular: No | Convergence: OK

**Table 6.** FDS MEPs LMM Fixed effects.

| Term | Estimate | SE | df | t | *p* | Sig. |
| --- | --- | --- | --- | --- | --- | --- |
| (Intercept) | 4.966 | 0.194 | 15.0 | 25.60 | < 0.001 | *** |
| Pred. Unpredictable | 0.068 | 0.023 | 41.2 | 2.93 | 0.005 | ** |
| Error_Prev Failure | 0.018 | 0.019 | 115.6 | 0.96 | 0.338 |  |
| Block_Factor4 | 0.048 | 0.037 | 17.9 | 1.30 | 0.210 |  |
| Block_Factor6 | 0.154 | 0.048 | 16.6 | 3.21 | 0.005 | ** |
| trial_in_block_z | 0.028 | 0.015 | 15.0 | 1.83 | 0.088 | . |
| Pred×Error | −0.023 | 0.025 | 9270 | −0.92 | 0.358 |  |
| Pred×Block4 | 0.024 | 0.023 | 9249 | 1.05 | 0.295 |  |
| Pred×Block6 | 0.018 | 0.023 | 9251 | 0.80 | 0.422 |  |
| Error×Block4 | 0.032 | 0.025 | 9285 | 1.32 | 0.188 |  |
| Error×Block6 | −0.016 | 0.025 | 9283 | −0.66 | 0.510 |  |
| Pred×Error×Block4 | −0.008 | 0.036 | 9265 | −0.22 | 0.822 |  |
| Pred×Error×Block6 | 0.002 | 0.036 | 9261 | 0.06 | 0.949 |  |

*Note.* Significance codes: \*\*\* $p < 0.001$ , \*\* $p < 0.01$ , \* $p < 0.05$ , . $p < 0.10$ .

**Table 7.** Type III ANOVA (Satterthwaite) + Effect Size.

| Term | F | NumDF | DenDF | $p$ | $\eta_p^2$ |
| --- | --- | --- | --- | --- | --- |
| Predict | 14.79 | 1 | 15.3 | 0.002** | 0.49 |
| Prev Error | 0.99 | 1 | 15.0 | 0.335 | 0.06 |
| Block | 7.54 | 2 | 15.2 | 0.005** | 0.50 |
| Trial within Block | 3.34 | 1 | 15.0 | 0.088 | 0.18 |
| Predict × Prev Error | 2.84 | 1 | 9269.7 | 0.092 | $< 0.001$ |
| Predict × Block | 0.81 | 2 | 9250.9 | 0.444 | $< 0.001^{**}$ |
| Prev Error × Block | 2.96 | 2 | 9270.0 | 0.052 | $< 0.001$ |
| Predict × Prev Error × Block | 0.04 | 2 | 9261.5 | 0.956 | $< 0.001^{**}$ |

*Note.* F = F statistic; NumDF = numerator degrees of freedom; DenDF = denominator degrees of freedom; $p$ = p-value; $\eta_p^2$ = partial eta squared. Trial within Block was z-scored. Asterisks indicate statistically significant effects: $p < 0.05$ (\*), $p < 0.01$ (\*\*), and $p < 0.001$ (\*\*\*).

**Table 8.** Variance Components (random effects)

| Component | Std.Dev. |
| --- | --- |
| Intercept (volunteer) | 0.7750** |
| P_unpred | 0.0665 |
| E_error | 0.0351 |
| B4 | 0.1373 |
| B6 | 0.1838** |
| trial_in_block_z | 0.0588** |
| Residual | 0.3339 |

<sup>38</sup> **Table 9** *Model Fit ( $R^2$ )*

**Table 9.** Model Fit (R*)

| Metric | Value |
| --- | --- |
| Conditional $R^2$ | 0.009 |
| Marginal $R^2$ | 0.845 |

<sup>39</sup> **Table 10** *FDS MEPs analysis - Model's comparison table*

**Table 10.** FDS MEPs analysis Model’s comparison table.

| Term | Model | F | $p$ | $\eta_p^2$ |
| --- | --- | --- | --- | --- |
| Predictability | Max | 14.794 | 0.0015 | 0.492 |
|  | RMA | 13.747 | 0.002 | 0.478 |
| Previous error | Max | 0.990 | 0.3355 | 0.062 |
|  | RMA | 0.923 | 0.352 | 0.058 |
| Block | Max | 7.540 | 0.0053 | 0.498 |
|  | RMA | 6.912 | 0.005 | 0.315 |
| Trial within<br>block | Max | 3.335 | 0.0878 | 0.182 |
|  | RMA | — | — | — |
| Predictability ×<br>Previous error | Max | 2.843 | 0.0918 | 0.000 |
|  | RMA | 4.554 | 0.050 | 0.233 |
| Predictability ×<br>Block | Max | 0.812 | 0.4440 | 0.000 |
|  | RMA | 0.754 | 0.450 | 0.048 |
| Previous error ×<br>Block | Max | 2.963 | 0.0517 | 0.001 |
|  | RMA | 1.689 | 0.208 | 0.101 |
| Predictability ×<br>Previous error ×<br>Block | Max | 0.045 | 0.9565 | 0.000 |
|  | RMA | 0.055 | 0.943 | 0.004 |

*Note.* Max = maximal model; RMA = repeated-measures ANOVA; $\eta_p^2$ = partial eta squared.
Dashes indicate that the effect was not included in the RMA model.

#### FDS Cell-aggregate RM-ANOVA

The aggregated RM-ANOVA confirmed the main effects of Predictability (RM-ANOVA: F(1, 15) = 13.75, *p* = 0.002, 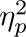 = 0.478) and (F(1.78, 26.73) = 6.91, *p* = 0.005, 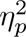 = 0.315) Block obtained in LMM. The two analyses diverged only on the weaker Predictability × Previous-Error term, which was significant in the aggregated RM-ANOVA (F(1, 15) = 4.55, *p* = 0.050, 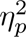 = 0.233). Given its inconsistency across specifications, we do not interpret it further. Both confirmed the main effects of Predictability (RM-ANOVA: F(1, 15) = 13.75, *p* = 0.002, 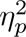 = 0.478) and Block (F(1.78, 26.73) = 6.91, *p* = 0.005, 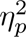 = 0.315), with no significant three-way interaction (F(1.94, 29.13) = 0.06, *p* = 0.943). The two analyses diverged only on the weaker Predictability × Previous-Error term, which was non-significant in the trial-level models (robust *p* = 0.092; intercept-only *p* = 0.174) but reached the threshold in the aggregated RM-ANOVA (F(1, 15) = 4.55, *p* = 0.050, 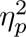 = 0.233).

#### Linear Mixed-Effects Model Predicting log(RT)

Formula:

log_RT ∼ Predictability * Error_Prev * Block_Factor (1 | volunteer)

Obs: 9221 | Groups: 16 volunteers | REML: 9520.1 | Singular: No | Convergence: OK

**Table 11.**
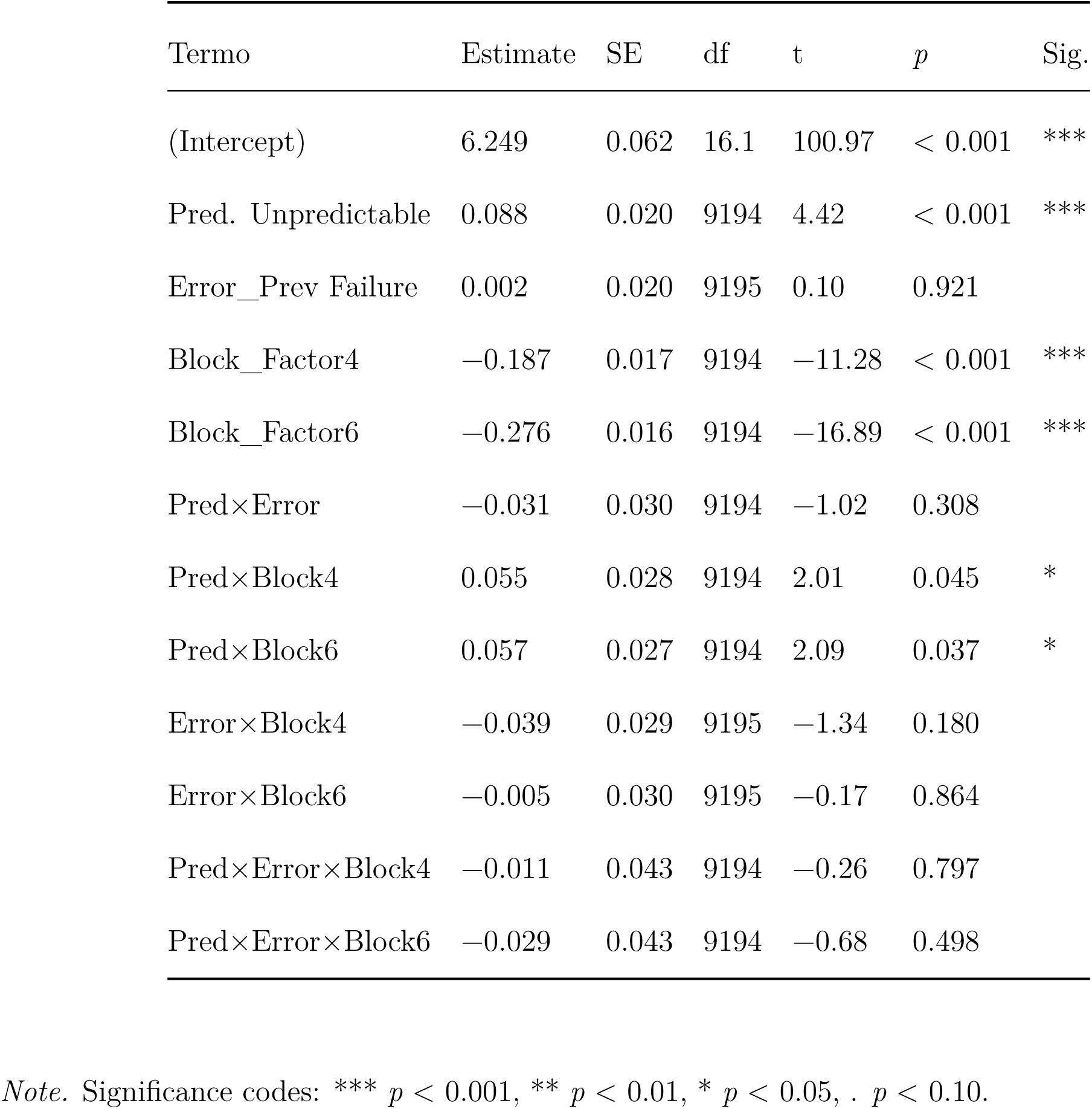
RTs MEPs LMM Fixed effects.

| Termo | Estimate | SE | df | t | $p$ | Sig. |
| --- | --- | --- | --- | --- | --- | --- |
| (Intercept) | 6.249 | 0.062 | 16.1 | 100.97 | $< 0.001$ | *** |
| Pred. Unpredictable | 0.088 | 0.020 | 9194 | 4.42 | $< 0.001$ | *** |
| Error_Prev Failure | 0.002 | 0.020 | 9195 | 0.10 | 0.921 |  |
| Block_Factor4 | -0.187 | 0.017 | 9194 | -11.28 | $< 0.001$ | *** |
| Block_Factor6 | -0.276 | 0.016 | 9194 | -16.89 | $< 0.001$ | *** |
| Pred×Error | -0.031 | 0.030 | 9194 | -1.02 | 0.308 |  |
| Pred×Block4 | 0.055 | 0.028 | 9194 | 2.01 | 0.045 | * |
| Pred×Block6 | 0.057 | 0.027 | 9194 | 2.09 | 0.037 | * |
| Error×Block4 | -0.039 | 0.029 | 9195 | -1.34 | 0.180 |  |
| Error×Block6 | -0.005 | 0.030 | 9195 | -0.17 | 0.864 |  |
| Pred×Error×Block4 | -0.011 | 0.043 | 9194 | -0.26 | 0.797 |  |
| Pred×Error×Block6 | -0.029 | 0.043 | 9194 | -0.68 | 0.498 |  |

*Note.* Significance codes: \*\*\* $p < 0.001$ , \*\* $p < 0.01$ , \* $p < 0.05$ , . $p < 0.10$ .

**Table 12.** Type III ANOVA (Satterthwaite) + Effect Size.

| Term | F | NumDF | DenDF | $p$ | $\eta_p^2$ |
| --- | --- | --- | --- | --- | --- |
| Predict | 137.05 | 1 | 9194.1 | < 0.001*** | 0.01 |
| Prev Error | 15.51 | 1 | 9194.9 | < 0.001*** | < 0.01 |
| Block | 301.40 | 2 | 9194.1 | < 0.001*** | 0.06 |
| Predict × Prev Error | 6.27 | 1 | 9194.3 | 0.012* | < 0.01 |
| Predict × Block | 3.14 | 2 | 9194.0 | 0.043* | < 0.01 |
| Prev Error × Block | 2.18 | 2 | 9194.6 | 0.113 | < 0.01 |
| Predict × Prev Error × Block | 0.23 | 2 | 9194.1 | 0.792 | < 0.01 |

*Note.* F = F statistic; NumDF = numerator degrees of freedom; DenDF = denominator degrees of freedom; $p$ = $p$ -value; $\eta_p^2$ = partial eta squared. Asterisks indicate statistically significant effects: $p < 0.05$ (\*), $p < 0.01$ (\*\*), and $p < 0.001$ (\*\*\*).

**Table 13.** Variance Components (random effects)

| Component | Standard Deviation |
| --- | --- |
| Participant (Intercept) | 0.2428 |
| Residual | 0.4022 |

**Table 14.** Model Fit (R*)

| Metric | Value |
| --- | --- |
| Marginal $R^2$ | 0.063 |
| Conditional R<sup>2</sup> | 0.313 |

**Table 15.** RTs analysis Model’s comparison table.

| Term | Model | F | $p$ | $\eta_p^2$ |
| --- | --- | --- | --- | --- |
| Predictability | Par | 137.054 | < 0.001 | 0.015 |
|  | RMA | 29.028 | < 0.001 | 0.659 |
| Previous Error | Par | 15.509 | < 0.001 | 0.002 |
|  | RMA | 18.436 | < 0.001 | 0.551 |
| Block | Par | 301.401 | < 0.001 | 0.062 |
|  | RMA | 33.604 | < 0.001 | 0.691 |
| Predict × Prev Error | Par | 6.271 | 0.012 | 0.001 |
|  | RMA | 7.449 | 0.016 | 0.332 |
| Predict × Block | Par | 3.139 | 0.043 | 0.001 |
|  | RMA | 1.388 | 0.265 | 0.085 |
| Prev Error × Block | Par | 2.184 | 0.113 | < 0.001 |
|  | RMA | 1.552 | 0.230 | 0.094 |
| Predict × Prev Error × Block | Par | 0.233 | 0.792 | < 0.001 |
|  | RMA | 0.729 | 0.483 | 0.046 |

*Note.* Par = parsimonious; RMA = repeated-measures ANOVA; $\eta_p^2$ = partial eta squared. Dashes
indicate that the effect was not included in the RMA model.

Log-transformed RTs were analyzed with linear mixed-effects models including the maximal Predictability × Previous-Error × Block interaction. We first attempted to fit a maximal-by-design structure with uncorrelated random intercepts and slopes per volunteer (for Predictability, Previous-Error, Block, and trial position), matching the approach used for the MEP analysis. However, this model produced a singular fit, indicating that the data did not support the maximal random-effects structure. We therefore report the parsimonious random-intercepts model (random intercept per volunteer), which converged normally and was not singular (9221 observations, 16 participants).

RTs showed three main effects together with a theoretically central interaction. RTs were faster on predictable than unpredictable trials (F(1, 9194) = 137.05, *p* < 0.001, 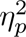 = 0.01; ≈ 441 vs. ≈ 489 ms) and became faster across the game, dropping gradually from Block 2 to Block 6 (F(2, 9194) = 301.40, *p* < 0.001, 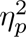 = 0.06; 537 → 415 ms; Block 6 / Block 2 ratio = 0.77, z = −23.79, *p* < 0.001, Fig. S1). A Previous-Error main effect was also present (F(1, 9195) = 15.51, *p* < 0.001, 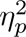 < 0.01), and Predictability interacted with the Previous-Error (F(1, 9194) = 6.27, *p* = 0.012, 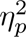 < 0.01). Decomposing this interaction, the slowing cost of unpredictability was larger after a success (predictable/unpredictable ratio = 0.882, z = −11.31, *p* < 0.001) than after a failure (ratio = 0.922, z = −5.91, *p* < 0.001); read the other way, the prior outcome mattered only under uncertainty, where participants were faster on the next unpredictable trial following a failure (success/failure ratio = 1.06, z = 4.45, *p* < 0.001), whereas on predictable trials it had no effect (ratio = 1.01, *p* = 0.294). In short, a previous failure selectively sharpened the RT when the upcoming context was uncertain. A weaker Predictability × Block interaction reached significance (F(2, 9194) = 3.14, *p* = 0.043, 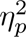 < 0.01), but simple-effects analyses showed the predictability cost was present in every block (all *p* ≤ 0.001) and varied only marginally in magnitude; we therefore treat it as a minor, scale-dependent modulation rather than a robust effect. The remaining interactions, including the three-way term, were non-significant (three-way: *p* = 0.792, 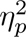 < 0.01). As for the MEPs, a sensitivity RM-ANOVA on trial-averaged data confirmed the main effects of Predictability and Previous-Error and the Predictability × Previous-Error interaction (Supplementary Results: Table 15). Because the trial-level model was intercepts-only for these terms, its very large F values (137.05 and 301.40) are anticonservative; the reliable effect magnitudes are those from the RM-ANOVA (F = 29.03 and F = 33.60).

#### RT Cell-aggregate RM-ANOVA

To confirm that these conclusions did not depend on the trial-level modeling approach, we crossvalidated all effects against an aggregated repeated-measures ANOVA computed on participant-level cell means (Greenhouse–Geisser corrected). This analysis showed significant main effects of Predictability (F(1, 15) = 29.03, *p* < 0.001, 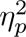 = 0.66), Block (F(1.68, 25.19) = 33.60, *p* < 0.001, 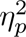 = 0.69), and Previous-Error (F(1, 15) = 18.44, *p* < 0.001, 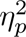 = 0.55), and a significant Predictability × Previous-Error interaction (F(1, 15) = 7.45, *p* = 0.016, 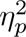 = 0.33). The Predictability × Block interaction was not significant in the aggregated analysis (F(1.84, 27.57) = 1.39, *p* = 0.265), reinforcing our cautious interpretation of that term, and the three-way interaction remained null (*p* = 0.483).

## Supplementary Figures

**Figure S1:**
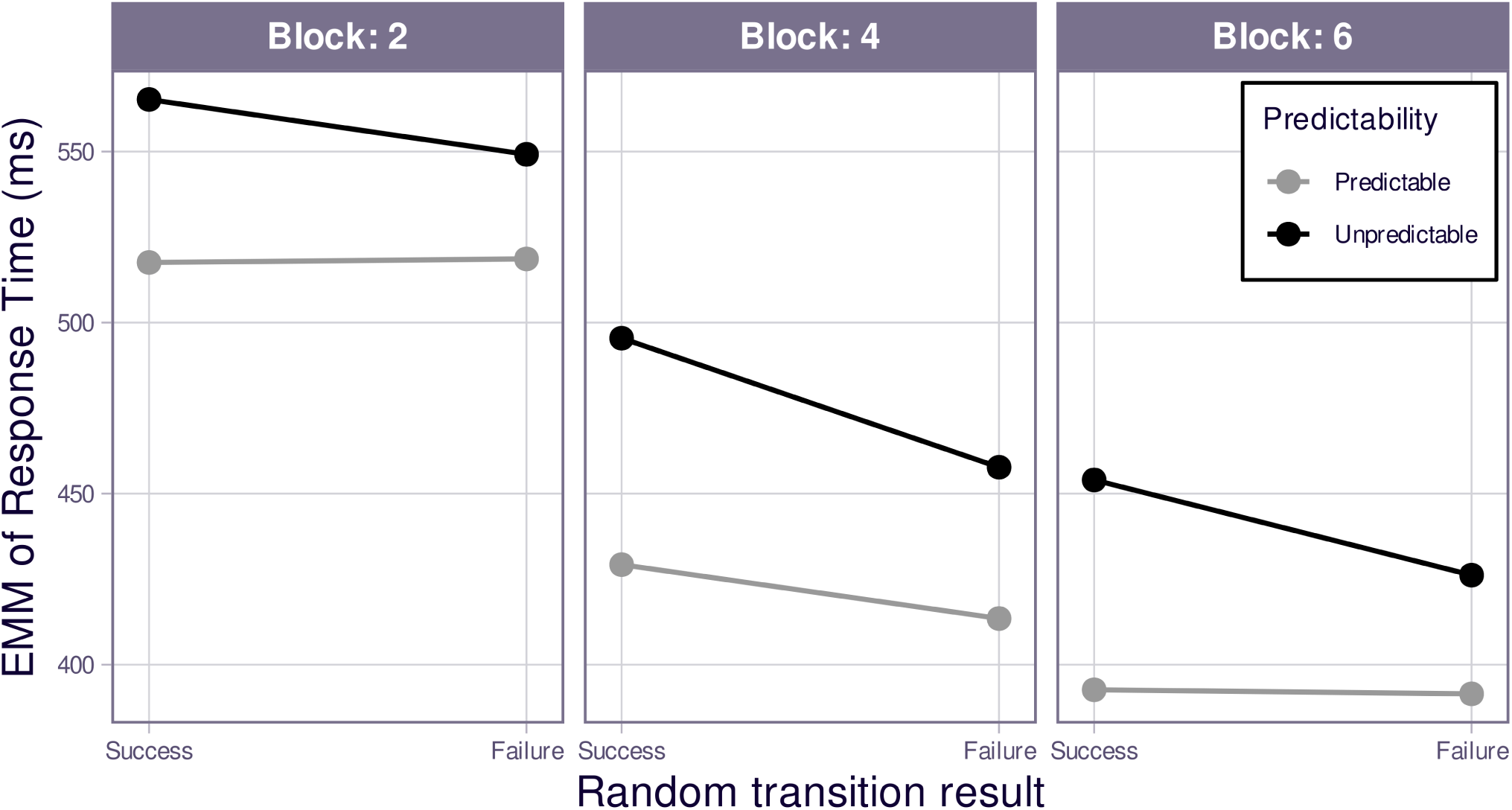
Effects of block (2, 4, 6), previous prediction outcome in non-deterministic transitions (failure, success), and contextual predictability (predictable, unpredictable) on Response Times (RTs). The y-axis shows estimated marginal means (EMMs) of the RTs derived from the linear mixed-effects model (LMM). The x-axis is organized in two hierarchical levels: block at the outer level (Block 2, Block 4, Block 6) and previous prediction outcome at the inner level (failure, success). Line colors represent context predictability (gray: predictable; black: unpredictable). *n* = 16.

**Figure S2:**
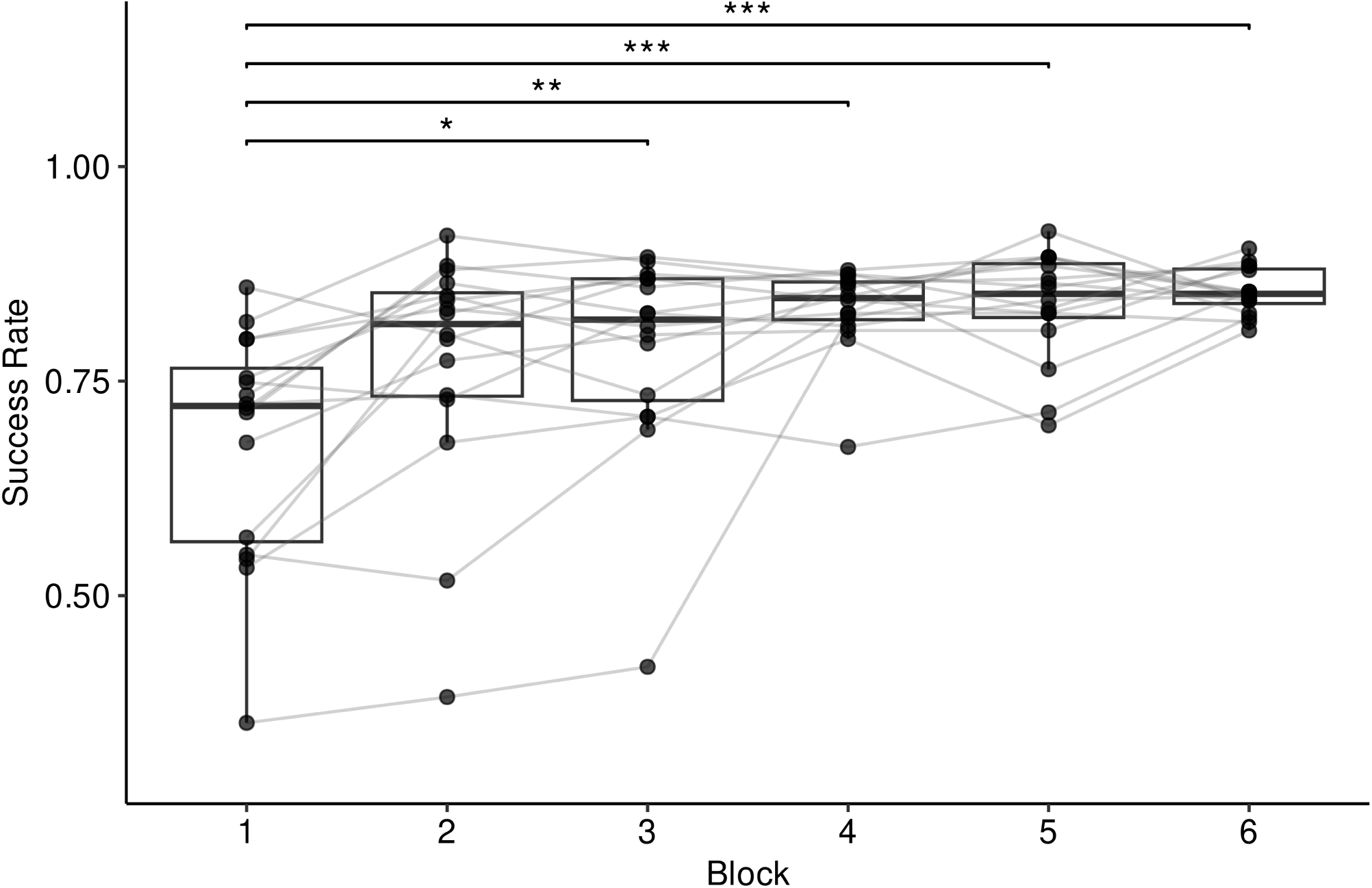
Progression of success rate across blocks in the Goalkeeper Game. The x-axis represents the game blocks and the y-axis indicates the success rate. *n* = 16.

